# Evaluation of Cross-slice Imputation for Spatial Proteomics

**DOI:** 10.64898/2026.10.27.762306

**Authors:** Georgiy A. Bondar, Behnaz Bozorgui, Anil Korkut, Abhishek Halder

**Affiliations:** Department of Applied Mathematics, University of California Santa Cruz, USA; Department of Bioinformatics and Computational Biology, UT MD Anderson Cancer Center, Houston, TX 77030, USA; Department of Aerospace Engineering, Iowa State University, USA

## Abstract

Spatial single-cell protein profiling is at the forefront of translational medicine research efforts that try to decipher tissue heterogeneity. However, existing experimental approaches are limited in the number of biomarkers that they measure. This has motivated extensive research into the problem of protein imputation, where proteins or biomarkers not directly measured in a given slice of biopsy tissue are learned from those measured.

In this work, we focus on the specific problem of cross–slice protein imputation (CSPI). In the CSPI problem, two sets of biomarker expressions at the single-cell level are known for physically adjacent slices of tissue. Each slice has its own known set of biomarkers, which overlaps with but does not subsume that of the adjacent. For each slice, we wish to learn the expressions of the proteins unique to the adjacent.

Existing literature for protein imputation methodologies can be grouped in two categories: learning-based and alignment-based approaches. Learning-based approaches do not account for spatial and structural similarities between the training and the target sets. Alignment-based approaches aim to determine an association between cells in adjacent slices, but without the explicit goal of using this alignment for imputation. In this work, we evaluate several methods from both categories and their adaptibility to the cross-slice protein imputation problem. We provide a statistical analysis of all methods’ performance under varying conditions of implementation. We find that learning-based methods significantly outperform alignment-based methods at the task of cross-slice protein imputation, with the latter more suited to the task of inter-slice interpolation or 3D tissue reconstruction. Moreover, inter-patient imputation is far less accurate than intra-patient imputation for all methods evaluated.

## 1 Introduction

Spatial omics methods have transformed our understanding of tissue structures in both diseased and healthy states (Elhanani et al.; Goltsev et al.; Lin et al., 2018; Moffitt et al.). Such methods can inform on key clinical and biological biomarkers such as proteins that identify cellular linages, cell states, or therapy responses. Spatial proteomics methodologies are particularly critical as proteomics-based markers are the backbone of pre-clinical and clinical diagnostic assays such as immunohistochemistry (Bollhagen & Bodenmiller; Galon et al.; Keren et al.). Existing spatial proteomics methods benefit form highly multiplexed imaging of tissue slices stained with fluorescently labeled antibodies (Giesen et al.; Lin et al., 2018). However, two major impediments limit spatial proteomics. First, the current multiplexed imaging based spatial proteomics techniques are limited to 40-50 proteins per tissue slice that prevents comprehensive molecular coverage (Bollhagen & Bodenmiller). Second, imaging is limited to 5 micron thick slices, providing 2-dimensional information on a 3-dimensional tissue. Solution of these two major challenges has the potential to enable readouts for a larger set of proteins, leading to insight on diverse tissue phenotypes and assessment of tissue characteristics in 3-dimensions across the continuum of conventional tissue sections.

Several computational solutions have been proposed to address these major challenges. These methods infer the expression of specific biomarkers from the known expressions of others through a variety of imputation methods across multiple tissue sections. For instance, the work by Kirchgaessner et al. (2025) deploys machine learning techniques such as regression, gradient-boosted trees, and autoencoders to impute missing or corrupted protein expression values at a single-cell level in multiplex tissue imaging (MTI) datasets. Alternatively, Heussner et al. (2025) proposes a method for integrating physically adjacent MTI images at a single-cell level, allowing for protein expression transfer between these aligned images. These two methods are representative of the two categories that most cross-slice protein imputation methods fall into – learning-based and alignment-based. Methods in this first category train a model on known spatial protein expression and tissue neighborhood patterns to predict unseen spatial expression profiles. Methods in the second category find a geometric alignment between spatially close tissue slices, forming associations between cells across slices – missing protein expression values can then be transferred between slices along these associations. Although both methodological categories have shown promise, their relative strengths and limitations remain poorly understood. In particular, there has not been a systematic evaluation of learning-based and alignment-based approaches for cross-section protein imputation. Such an assessment is essential for guiding method development and identifying the conditions under which different computational strategies could be preferred over others. The purpose of this work is to fill this research gap.

Here, we have performed a comprehensive benchmark of widely used spatially-resolved protein expression imputation methods, including MTI (Kirchgaessner et al., 2025), SevenUp (Wu et al., 2023), MIM-CyCIF (Sims et al., 2024), COEXIST (Heussner et al., 2025), and MOSCOT (Klein et al., 2023) (see Table 1). The former three are learning-based, and provide a broad survey of various machine learning architectures applied to the protein imputation problem. The latter two – alignment-based – offer two differing approaches of matching cells across slices. To achieve an objective comparison between these methods, we make use of the `CRC_atlas_2022` dataset (Lin et al., 2023), which offers serial tissue sections from a colorectal cancer patient that were stained with overlapping protein marker sets. In each application, we used a set of protein markers on a given slice to predict protein marker expression with spatial resolution in an adjacent tissue. This design of experiment enables objective evaluation of computational approaches by providing ground-truth protein measurements that can be withheld and subsequently predicted from adjacent sections. Specifically, we assess the ability of individual methods to infer protein expression in one tissue section using information from neighboring sections.

**Table 1:**
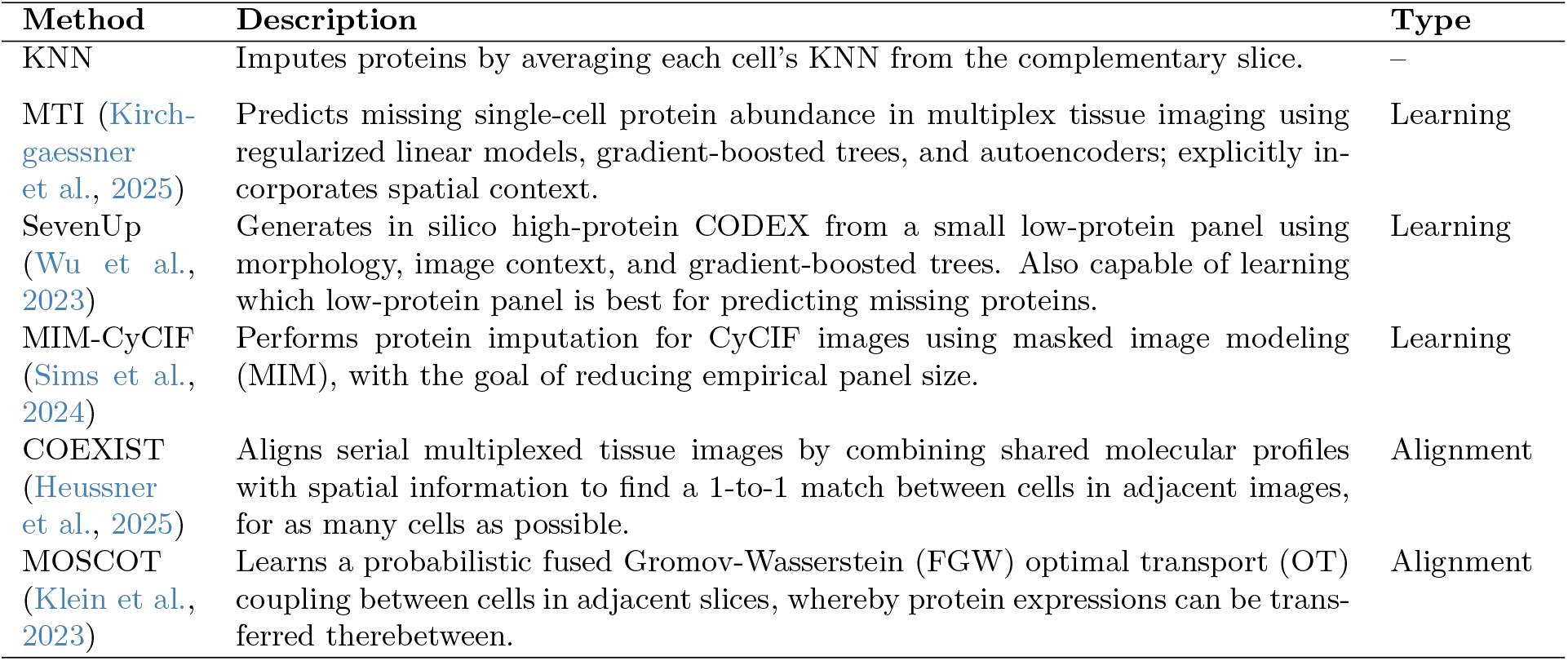
Existing spatial proteomics methods adaptable to the CSPI problem.

In this work, we are interested in the problem of *cross–slice* protein imputation (CSPI). Spatial proteomics experiments are limited in their imaging throughput – when a biopsy is performed, physically adjacent slices of tissue are oft extracted from the patient, giving several samples which can be dyed and bleached to obtain biomarker expressions therefor. It is the choice of the pathologist which biomarkers to dye for in each slice, up to a physical limit before cells are damaged and unusable. Thus, the total set of desired biomarkers can be split between two adjacent slices, and protein imputation methods can be used to reconstruct the protein measurements missing in each slice from those of the adjacent. To evaluate existing methods on this problem, we considered pairs of adjacent tissue sections containing a set of shared proteins and a set of proteins unique to each section. We assessed how accurately different methods could predict the proteins unique to one section using information available in its neighboring section. We formalize this in Problem 1 in Section 2.

Specifically, we evaluated the performance of several methods from both the learning and alignment-based classes to address the following questions:

1. How do the known protein sets in each slice affect the imputation performance across methods?
2. How does choosing slices further apart in space affect imputation performance across methods?

Overall, we found that learning-based methods significantly outperform alignment-based methods. Moreover, the MIM-CyCIF (Sims et al., 2024) and 7-UP (or SevenUp) (Wu et al., 2023) methods, both designed for protein panel upsampling, consistently performed the best across all benchmarks in this study.

This study will inform the development of new methods that for improved solution of the CSPI problem. The solution will help diversify tissue profiling in 2 and 3 dimensions. This will offer a deeper insight into proteomic composition of tissues and development of new precision therapy strategies such as those that target tumor-immune microenvironment in colorectal cancer (Schürch et al.).

### Outline

Section 2 describes the methods we evaluate, the dataset we work with, as well as their adaptation to our specific CSPI problem. We also discuss our benchmarking pipeline, which consists of two main benchmarks to evaluate the methods’ performance under varying conditions, as well as our process of evaluating the results thereof.

Section 3 presents the detailed results of our benchmarks, wherein we vary the data availability to the CSPI solution methods and the tissue slices. These benchmarks quantify the performance of each method, as well as across methods.

Section 4 provides a discussion of our benchmarking results of the previous section. We analyze the quantitative results of both benchmarks as well as the general observed patterns of method behavior and accuracy. We conclude the work with suggestions for future exploration of the CSPI problem, as well as with practical recommendations for application of existing methods therefor.

### Notations

Some standard notations used throughout are: | · |for the cardinality of a set, ⟬· ⟭ for a finite set, ⊙for elementwise multiplication, and ⊘ for elementwise division. The exponential and logarithm operators, denoted as exp and log respectively, are understood elementwise. The remaining notations are explained in situ.

## 2 Methods

In this work, we evaluate several existing protein imputation methods’ ability to solve the CSPI problem. Each of these methods will be run on the same data through our benchmarking pipeline, and its results compared with the others. This section provides the details of the methods, the dataset, the pipeline, and the evaluation scheme.

Figure 1 provides a visual overview of our dataset, methods, and the imputation methcanisms of learning vs. alignment-based methods. We begin by formalizing the CSPI problem.

**Figure 1:**
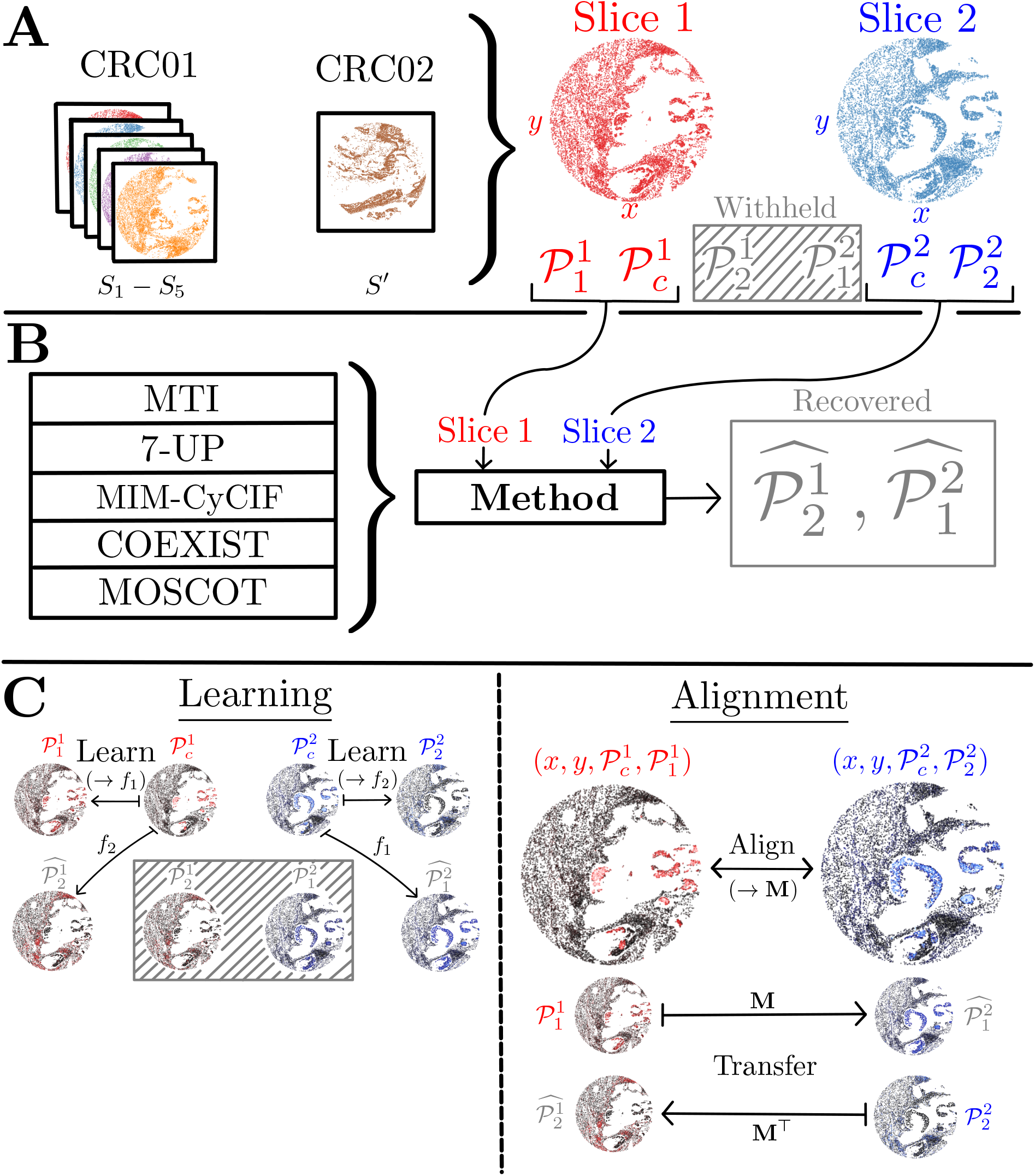
**A**. Our dataset consists of five slices *S*_1_ through *S*_5_ from CRC01 along with one slice *S*^′^ from CRC02. For each benchmark, we select two slices from this set. Each slice *i* has spatial features (*x, y*) and single-cell protein expressions 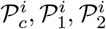. **B**. Our CSPI benchmarking pipeline consists of the selection of an imputation method, wherefrom 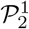 and 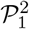 are withheld. Using the remaining features, the method recovers these withheld values via the respective predictions 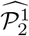 and 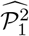. **C**. Methods we evaluate are either learning-based (MTI, SevenUp, MIM-CyCIF) or alignment-based (COEXIST, MOSCOT). A method of the former type will learn a function *f*_1_ from 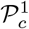 to 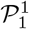 , which applied to 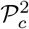 yields the prediction 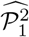 , and vice versa. An alignment-based method will instead learn an alignment (shown matricially as **M**) between known features of both slices. Features known uniquely to one slice can then be transferred along the alignment to the other slice, recovering the withheld features.

### Problem 1

(Single-cell Cross-slice Protein Imputation (CSPI)). *Consider a pair of 2-dimensional ‘slices’ of tissue – slice 1 and slice 2 – close together in the z-axis, orthogonal to the* (*x, y*) *slice plane. Each slice consists of n*_1_ *and n*_2_ *cells respectively, with each cell having a position* (*x, y*) *as well as a vector of protein expressions. The proteins expressed are* not *the same in both slices. Specifically, we denote by*

- *P*_*c*_ *the set of proteins common to both slices;*
- *P*_1_ *the set of proteins unique to slice 1;*
- *P*_2_ *the set of proteins unique to slice 2*.

*Thus the full protein set of interest is P*_1_ ⊔ *P*_*c*_ ⊔ *P*_2_ =: *P, where* ⊔ *denotes the disjoint union*.

*We desire to know expressions of all proteins P in each cell, in both slices. I*.*e*., *we wish to learn expressions of P*_2_ *for all cells in slice 1, and the same of P*_1_ *for all cells in slice 2*.

The proteins ‘unique’ to one slice are actually known in the other slice, but are withheld for the purpose of validation. We adopt the convention that 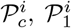, and 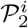 are the actual protein expressions for all cells in slice *i* ∈ *{*1 ,2 }, with 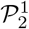 withheld for slice 1 and similarly 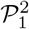 withheld for slice 2^1^.

### 2.1 Dataset

We will use the CRC_atlas_2022 dataset (Lin et al., 2023). This dataset consists of 25 slices of ≈400, 000 cells each, with 24 protein expressions per slice, all of which are common and given as the ‘full’ protein set

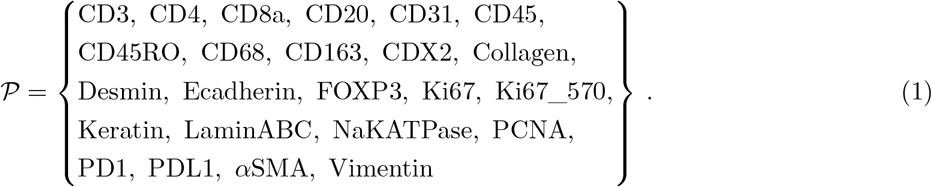

Thus, the mismatch in known proteins present in Problem 1 must be introduced methodically in the benchmarks. We use the tabular form of the dataset, meaning that each cell is assigned a numerical vector containing its position, morphology, and biomarker expressions.

For our benchmarks, we need several slices of tissue which are close together in space, as well a one which is far from the others. Moreover, we want to evaluate method performance with a smaller and larger amount of cells per slice. Thus motivated, we use six particular slices of tissue from the dataset in our benchmarks – Figure 2 shows the slices we use, each with ≈400 000 cells, as well as 10 000 cell slices we obtain by taking a localized section of cells from each.

**Figure 2:**
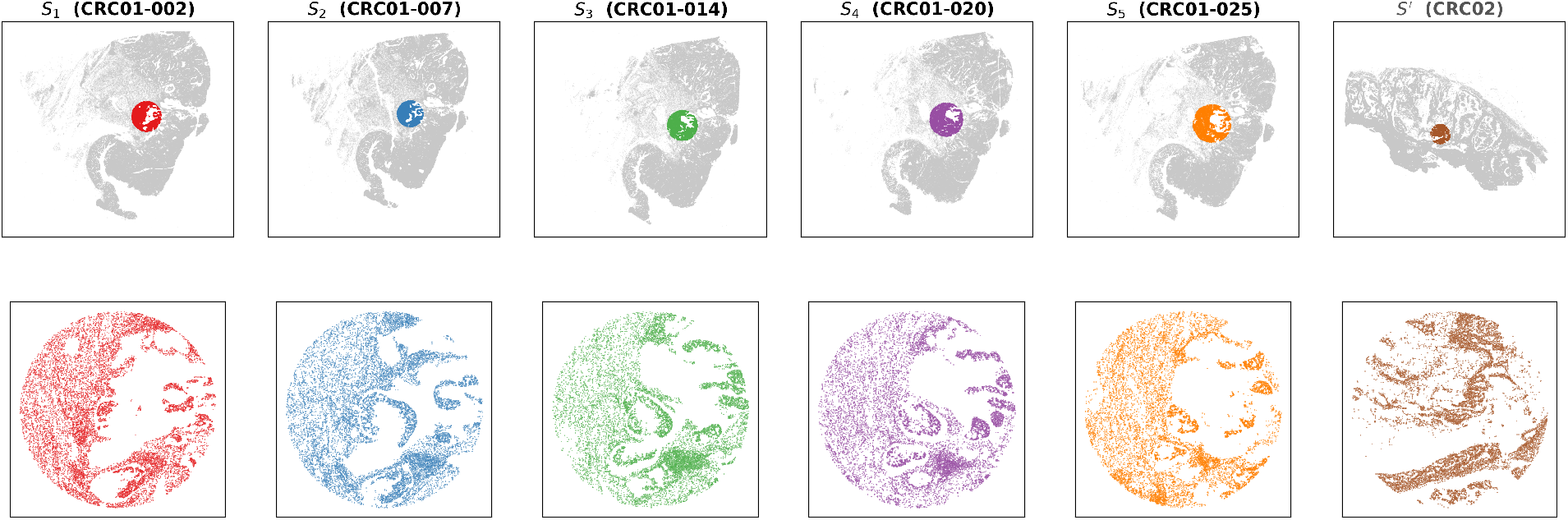
Slices *S*_1_ through *S*_5_ from patient CRC01, and *S*^′^ from patient CRC02. *S*_1_ and *S*_2_ are used in Benchmark 1, and all slices are used in Benchmark 2. The full slices are shown in the top row, with 10 000-cell sub-slices on the bottom row – the latter are obtained by taking the closest 10 000 cells from the same spatial reference point across slices.

Table 2 shows how we label the slices throughout this work, as well as the slices’ relationship with each other. Slices *S*_1_ through *S*_5_ are spatially close and from the same patient, whereas *S*^′^ is from a different patient.

**Table 2:** Tissue sections used in this study. *S*_1_–*S*_5_ are consecutive sections from the same CRC01 patient (WD-76845); *S*^′^ is a single section from a second patient CRC02 and will be used for cross-patient evaluation.

| Label | Name | Depth ($\mu\text{m}$) |
| --- | --- | --- |
| $S_1$ | CRC01-002 | 2 |
| $S_2$ | CRC01-007 | 7 |
| $S_3$ | CRC01-014 | 14 |
| $S_4$ | CRC01-020 | 20 |
| $S_5$ | CRC01-025 | 25 |
| $S'$ | CRC02 | — |

### 2.2 Normalization

Raw protein expression values in the `CRC_atlas_2022` dataset are obtained from pixel intensity values in images of stained slices. These values are affected by variability of light exposure, either over an entire slice or locally within. Moreover, the process of tabulating these values at the single-cell level can result in artifacts where individual cells have extreme protein expression values, which skew the overall distribution of these values within a slice. To circumvent these issues, we normalize all protein expression values prior to our benchmarking. For this normalization, we use a two-step procedure explained next applied independently to each slice.

First, we apply the log1p transform (Harris et al., 2020) *P*^*i*^ ⟼ log(1 + *P*^*i*^) to both slices *i* ∈ *{*1, 2} , where *P*^*i*^ are all protein expressions (known and withheld) in slice *i*. This compresses the dynamic range of protein expressions, makes the floor of expression zero, and makes the distribution of expressions more symmetric, as the log dampens extreme values. Second, these transformed values are then z-scored per protein (Hickey et al., 2021) as per the following transformation applied to each row of *P*^*i*^:

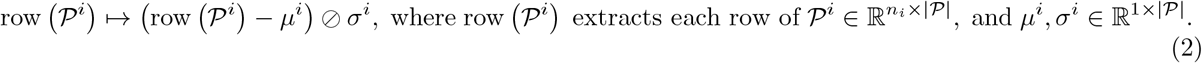

We note that these quantities are computed per-slice independently, i.e., each slice is scaled relative to its *own* expression distribution rather than a shared cross-slice reference. This choice avoids conflating measurable biological differences between slices with the normalization itself. All our tested methods fit and predict in this normalized space; predictions 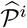 are denormalized before evaluation via the inverse two steps of normalization:

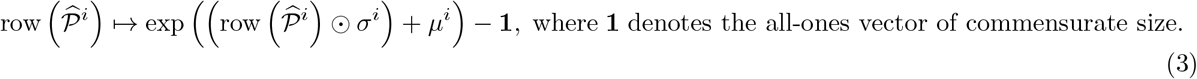

As before, row 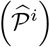 extracts each row of *P*^*i*^ and then applies the transformation (3). This ensures that Spearman *ρ* is computed on the original intensity scale. Non-protein columns (spatial coordinates, morphological features, Hoechst channels) are not normalized by the pipeline and are handled internally by each method as needed.

### 2.3 Methods Evaluated

Table 1 provides a summary of methods relevant to our Problem 1. We provide a more extensive description of each method below.

#### K-Nearest Neighbours (KNN)

This is the simplest among all of our evaluated methods and requires neither learning nor alignment. Instead, the cells of each slice are stored as a lookup index over a combined feature space of *P*_*c*_ expression and spatial coordinates (*x, y*), with each group independently z-scored and reweighted by a spatial or expression blending factor. For a fixed positive integer *k* = 15, each query cell from one slice finds its *k* nearest neighbors in the other slice’s index, and the imputed target protein value is the weighted average of those neighbors’ measured values. Two independent indices are built: one rooted in slice 2 (used to predict _2_ for slice 1) and one rooted in slice 1 (used to predict *P*_1_ for slice 2). In each case, the other slice is used strictly as a retrieval database — no parameters are estimated, and no information from the query slice influences the index. We include KNN in this work as a methodologically simple baseline for all other methods.

#### MTI (Kirchgaessner et al., 2025)

Originally developed for cross-panel protein imputation within single multiplexed tissue imaging datasets, three regression back-ends were proposed in Kirchgaessner et al. (2025): an `ElasticNet` linear model with cross-validated regularization, a `LightGBM` gradient-boosted tree ensemble, and a bottleneck autoencoder. We implement `ElasticNet` and `LightGBM`, each wrapped in a sklearn `MultiOutputRegressor`, with one model per target protein (Pedregosa et al., 2011). The feature vector for each cell is the concatenation of its own _*c*_ expression values and the spatially averaged *P*_*c*_ expression of all cells within a fixed radius (46 µm), capturing microenvironmental context. From this feature vector, the vector of expressions of proteins in the unknown set is learned. Our adaptation uses the same cross-slice training approach as all other methods: the reference slice provides both features and labels during training, the query slice provides features only during inference.

#### SevenUp (Wu et al., 2023)

This method was developed to computationally generate a 40–protein CODEX (co-detection by indexing) panel from a standard 7–protein Multiplex immunofluorescence (mIF) panel, motivated by the ubiquity of the latter data and practical limitations (time, cost) in obtaining the former in clinical settings. The method first uses a concrete autoencoder to learn, from full CODEX data, the seven biomarkers which, if known from the mIF panel, result in the best prediction from mIF to CODEX panels. A gradient-boosted tree algorithm is then trained to impute the remaining CODEX biomarkers from these seven. For benchmarks with fixed *P*_*c*_, the first part of this approach is not needed; indeed, in those cases it is essentially identical in approach to MTI. As with MTI above, this method trains solely on one slice to predict on the other.

#### MIM–CyCIF (Sims et al., 2024)

Masked Imaging Modeling for enhancing Cyclic ImmunoFluorescence (MIM-CyCIF) is a method for in-silico protein imputation in CyCIF data, designed to reduce panel acquisition cost by measuring only a subset of desired proteins empirically, and computationally recovering the rest. In the original formulation, a Vision Transformer (ViT)-based masked autoencoder operates on single-cell image tiles, where each cell is represented as a 32×32 pixel, 25-channel image. During training, a fixed proportion of protein channels are randomly masked, and a ViT encoder–decoder reconstructs the obscured channels from the observed ones.

As we use tabular single–cell measurements from the `CRC_atlas_2022` dataset rather than raw image tiles, we adapted MIM-CyCIF to operate on scalar features, drawing on the conceptual parallel between the original’s per-channel token structure and the token-per-feature paradigm of the Feature Tokenizer transformer (Gorishniy et al., 2021). In our adaptation, each scalar input — *P*_*c*_ protein expression values together with morphological covariates (AREA, CIRC) and nuclear staining intensities (Hoechst0–9) — is projected to a *d*-dimensional embedding via a learned linear transformation summed with a learned per-feature positional embedding. A transformer encoder (4 layers, 4 attention heads, *d* = 128) processes the full feature sequence, and a prepended classification token (Devlin et al., 2019) aggregates cross-feature information via self-attention before being passed to a two-layer MLP regression head (GELU activation) that jointly predicts all target proteins. We further reframe the task as supervised cross-slice regression rather than self-supervised channel reconstruction: two models are trained independently (one per imputation direction), each on one slice for 50-150 epochs using AdamW with cosine annealing (Loshchilov & Hutter, 2017; 2019), and applied to the held-out slice without observing it during training.

#### COEXIST (Heussner et al., 2025)

This method identifes cells which are shared across slices, and then finds a one-to-one mapping therebetween. In effect, this aligns the two slices, allowing for inter-slice feature transfer. In our implementation, the cost of matching cell *i* in the query slice to cell *j* in the reference slice is a blend of Pearson correlation over shared protein expression and normalized Euclidean distance in registered spatial coordinates, weighted equally. A greedy bijective matching is then solved to match ‘shared’ cells between slices. Cells that fail to find an acceptable match are handled by a KNN fallback trained on the matched reference cells. COEXIST is transductive: both slices are visible during fitting, and the imputed proteins for a query cell are simply the measured values thereof of its matched counterpart in the reference slice.

#### MOSCOT (Klein et al., 2023)

Multi-Omics Single-Cell Optimal Transport (MOSCOT) approaches cross-slice imputation as an optimal transport (OT) problem. Rather than learning a mapping function, it computes a probabilistic correspondence matrix 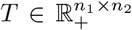 between all cells in slice 1 and 2 by solving a Fused Gromov-Wasserstein (FGW) problem. The FGW objective combines a linear term — the cost of moving probability mass between cells with dissimilar *P*_*c*_ expression vectors — with a Gromov-Wasserstein term — the cost of distorting the intra-slice spatial geometry to align the uncommon expression spaces *P*_1_ and *P*_2_. We use *α* = 0.3 (30% expression, 70% spatial), favouring cells that are spatially proximate and similarly positioned relative to their tissue neighbourhood. A rank-10 low-rank factorisation is used to make the 10 000 *×* 10 000 cell problem tractable on a single GPU. Once *T* is obtained, prediction is performed by matrix multiplication: 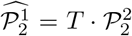, i.e., each slice 1 cell’s predicted expression is a weighted average of all slice 2 cells’ measured values, with weights given by the corresponding row of the transport plan. Like COEXIST, MOSCOT is transductive: the transport plan is solved jointly over both slices simultaneously, so both are visible during the “fitting” step and the result is specific to this pair of slices.

### 2.4 Method Benchmarking Strategy

Throughout this work, we run all methods to solve Problem 1 with the same full 24-protein set (1). Each run of a method requires the choice of method (given in Table 1), slices 1 and 2 (from Table 2), and the partitioning of features (*P*_*c*_, *P*_1_, *P*_2_).

All methods take as input the ‘known’ data for both slices, i.e, two tables containing per-cell spatial (*x, y*) data and protein expression (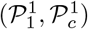 containing proteins *P*_1_⊔*P*_*c*_ for slice 1, and similarly 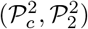 containing *P*_*c*_ ⊔ *P* _2_ for slice 2). Each method provides us with a prediction of the expressions of the withheld proteins for each slice, and for each cell within. We call the predictions for withheld proteins in slice 2 and slice 1 as 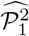 and 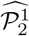, respectively.

Once predictions are made, we compare them to the known (and withheld) protein expressions for both slices – i.e., the withheld expressions 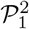 in slice 2 are compared with the predictions 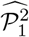. Therefor, we compute both Pearson (*r*) and Spearman (*ρ*) correlations between the withheld and predicted expressions, per-protein.

The rank-based Spearman correlation will be high if the ordering of cells by their expression for that protein is preserved by the method; i.e., if cells with relatively high withheld expression are predicted as such, and vice versa. Pearson correlation, however, is sensitive to the absolute magnitude of the predictions as well. Thus, high Spearman but lower Pearson correlation indicates that a method’s predictions are monotonically correct but skewed in magnitude.

### 2.5 Benchmark 1: Varying the protein coverage, |*P*_*c*_|

For this benchmark, we vary the number of proteins shared between slices. This allows us to quantify how protein imputation performance varies across methods. We test four different choices of *P*_*c*_, containing 4, 8, 12, and 16 proteins, as shown in Table 3. The core panel of four proteins was chosen to identify broad epithelial, immune, endothelial, and mesenchymal/stromal cell types. Subsequent proteins were added to enable finer discrimination of immune cell subsets (e.g., T cells, B cells, and macrophages), and to characterize functional cell states such as proliferation, and measure key immune checkpoint markers (PD-1 and PD-L1) with known predictive and prognostic significance in several cancer types. For each choice of *P*_*c*_, the remaining proteins are evenly and randomly (with fixed seed) split into *P*_1_ and *P*_2_. Throughout this benchmark, the choice of slices 1 and 2 remains fixed to *S*_1_ (`CRC01-002`) and *S*_2_ (`CRC01-007`) respectively (see Figure 2). I.e., we are using adjacent slices of tissue from the same patient, 5um apart. We perform first a scaled-down version of this benchmark where we use only a localized section of ≈ 10 000 cells in each slice, followed by using the full slices, each containing ≈ 400 000 cells.

**Table 3:** Shared-panel configurations used in Benchmark 1. Each set is a strict superset of the previous; proteins added relative to the preceding set are shown in bold.

| Set | $\vert \mathcal{P}_c\vert $ | Added proteins | Full $\mathcal{P}_c$ |
| --- | --- | --- | --- |
| 1 | 4 | — | Ecadherin, CD45, CD31, Vimentin |
| 2 | 8 | Keratin, PCNA, CD3, CD68 | Ecadherin, CD45, CD31, Vimentin, **Keratin, PCNA, CD3, CD68** |
| 3 | 12 | Ki67, CD20, α-SMA, CD163 | Ecadherin, CD45, CD31, Vimentin, Keratin, PCNA, CD3, CD68, **Ki67, CD20, α-SMA, CD163** |
| 4 | 16 | CD8a, CD4, PD1, PD-L1 | Ecadherin, CD45, CD31, Vimentin, Keratin, PCNA, CD3, CD68, Ki67, CD20, α-SMA, CD163, **CD8a, CD4, PD1, PD-L1** |

### 2.6 Benchmark 2: Varying inter-slice distance (*S*_1_ and *S*_2_)

For this benchmark, we fix a partition of *P* into *P*_*c*_, *P*_1_, *P*_2_ as set 2 from benchmark 1 (see Table 3) and solve the CSPI problem between different combinations of slices from the `CRC_atlas_2022` dataset. Specifically, we evaluate the performance of our methods on all combinations of slices from Table 2. As slices 1 and 2 must be different, and our methods’ performance should be symmetric, this gives us 15 possible unique combinations for our 6 slices. This benchmark allows us to gauge robustness of the methods’ performance to different slice pairs, different distances between slices, and for slices across two different patients.

## 3 Results

With the goal of evaluating several methods for the solution of the CSPI problem, our benchmarks 1 and 2 isolate the parameters of feature knowledge (i.e., how the protein set *P* is partitioned into *P*_1_, *P* _2_, and *P*_*c*_) and the choice of slice pairings (i.e., what is slice 1 and slice 2), respectively. In particular, the goal of benchmark 1 is to evaluate how methods perform when the number of commonly known proteins between slices (and as a consequence, the number to be imputed per-slice) is varied. The goal of benchmark 2 is to evaluate how varying the slices themselves affects performance.

Additionally, we run both benchmarks with 10 000-cell subslices, obtained by taking the closest 10 000 cells^2^ to an arbitrary reference point, and with the full slices (see Figure 2). This captures how the number of cells per slice can affect method performance, and allows us to efficiently run the alignment-based methods. These methods scale poorly to the full slices, which each contain ≈ 400 000 cells each^3^.

Below we report the results for both benchmarks.

### 3.1 The impact of protein coverage on imputation accuracy

Fixing slices 1 and 2 as *S*_1_ and *S*_2_ (see Table 2 and Figure 2) respectively, we ran all methods in Table 1 with the choices of *P*_*c*_, *P*_1_, and *P*_2_ outlined in Table 3 and Section 2.5.

The results are shown in Table 4 and Figure 3. Table 4 shows imputation accuracy of all methods in both Spearman and Pearson correlation coefficients, averaged over all target proteins. Figure 3 shows the same graphically (left), and also shows per-protein performance for all methods (right). We observe the expected behavior that all methods’ performance in both metrics increases with more common proteins, and that Pearson *r* remains consistently worse than Spearman *ρ*.

**Table 4:** Mean Spearman *ρ* / Pearson *r*, averaged over all target proteins and both imputation directions in Benchmark 1, as a function of the shared-panel size |*P*_*c*_|. The top table shows resuts for 10 000 cell-slices, and the bottom for full slices. The best results for each metric and |*P*_*c*_| size within each section are shown in **bold**.

| Method | $\vert \mathcal{P}_c\vert = 4$ | $\vert \mathcal{P}_c\vert = 8$ | $\vert \mathcal{P}_c\vert = 12$ | $\vert \mathcal{P}_c\vert = 16$ |
| --- | --- | --- | --- | --- |
| **10,000 cells per slice** |  |  |  |  |
| KNN | 0.6659 / 0.5845 | 0.7229 / 0.6251 | 0.7572 / 0.6820 | 0.7926 / 0.7356 |
| MTI-ElasticNet | 0.5913 / 0.5127 | 0.6724 / 0.5595 | 0.7179 / 0.6196 | 0.7551 / 0.6554 |
| MTI-LightGBM | 0.6656 / 0.5753 | 0.7462 / 0.6425 | **0.7790** / 0.6975 | 0.8299 / 0.7514 |
| SevenUp | 0.6772 / 0.5779 | 0.7455 / 0.6462 | 0.7735 / 0.6918 | 0.8308 / 0.7554 |
| COEXIST | 0.4269 / 0.3234 | 0.4822 / 0.3664 | 0.5164 / 0.4046 | 0.5621 / 0.4670 |
| MIM-CyCIF | **0.7013** / **0.5990** | **0.7564** / **0.6633** | 0.7771 / **0.6982** | **0.8343** / **0.7610** |
| MOSCOT | 0.4946 / 0.4474 | 0.5097 / 0.4322 | 0.5283 / 0.4633 | 0.5445 / 0.4807 |
| **Full slices** |  |  |  |  |
| KNN | 0.6791 / 0.5189 | 0.7258 / **0.5769** | 0.7391 / 0.6180 | 0.7562 / 0.6318 |
| MTI-ElasticNet | 0.6107 / 0.4210 | 0.6596 / 0.4999 | 0.6860 / 0.5396 | 0.7162 / 0.5544 |
| MTI-LightGBM | 0.6830 / 0.5072 | 0.7327 / 0.5743 | 0.7508 / 0.6119 | 0.7842 / 0.6591 |
| SevenUp | **0.7062** / **0.5258** | **0.7449** / 0.5755 | **0.7673** / 0.6233 | **0.8097** / 0.6747 |
| MIM-CyCIF | 0.6705 / 0.4997 | 0.7287 / 0.5687 | 0.7576 / **0.6285** | 0.8075 / **0.6900** |

**Figure 3:**
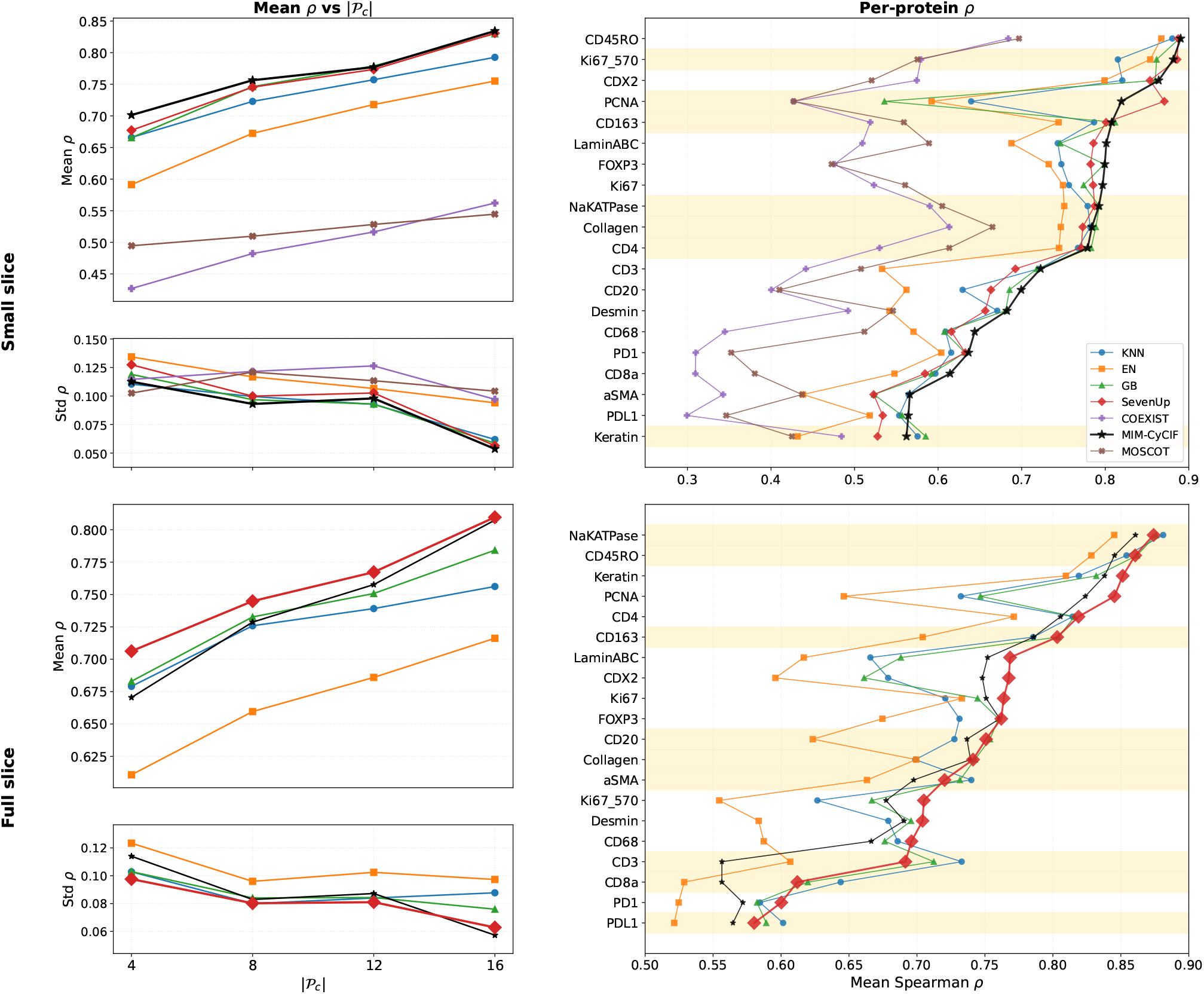
Spearman *ρ* of imputations from all methods in Benchmark 1, both averaged over all relevant *P* _*c*_ sets (left, along with standard deviations on bottom) and expressed per-protein (right). Highlighted proteins in the right are those where the average best-performing method is outperformed by at least one other method. For the small slice benchmark this best method is MIM-CyCIF; for the full slice benchmark it is SevenUp.

Notably, for the small-slice benchmark, MIM-CyCIF is the best performing method for most proteins, while SevenUp outperforms it on PCNA, and KNN with GB edge out MIM on Keratin. Worst-performing are both the alignment-based COEXIST and MOSCOT, with MTI’s ElasticNet performing the next-worst.

This performance trend, however, changes with the full-slice benchmark 1. With full slices, we see that SevenUp outperforms MIM-CyCIF on all proteins, and is significantly outperformed by other methods on CD3 only. For both small and full slice benchmarks, we see the standard deviation of predictions decrease as |*P*_*c*_| increases – this is both due to an immediately smaller pool of proteins to predict, as well as from real improvements in method precision with more training data.

### 3.2 Impact of inter slice distance on imutation accuracy

Fixing the choice of |*P*_*c*_| = 8 as in Table 3, we ran all methods in Table 1 between all possible 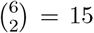 combinations of slices *S*_1_, … , *S*_5_, *S*^′^ as in Table 2. Figure 4 compares the *z*-axis distance between slices to imputation performance for all methods (top), and shows cross-slice imputation performance as mean Spearman *ρ* for all evaluated methods between all possible slice pairs (bottom).

**Figure 4:**
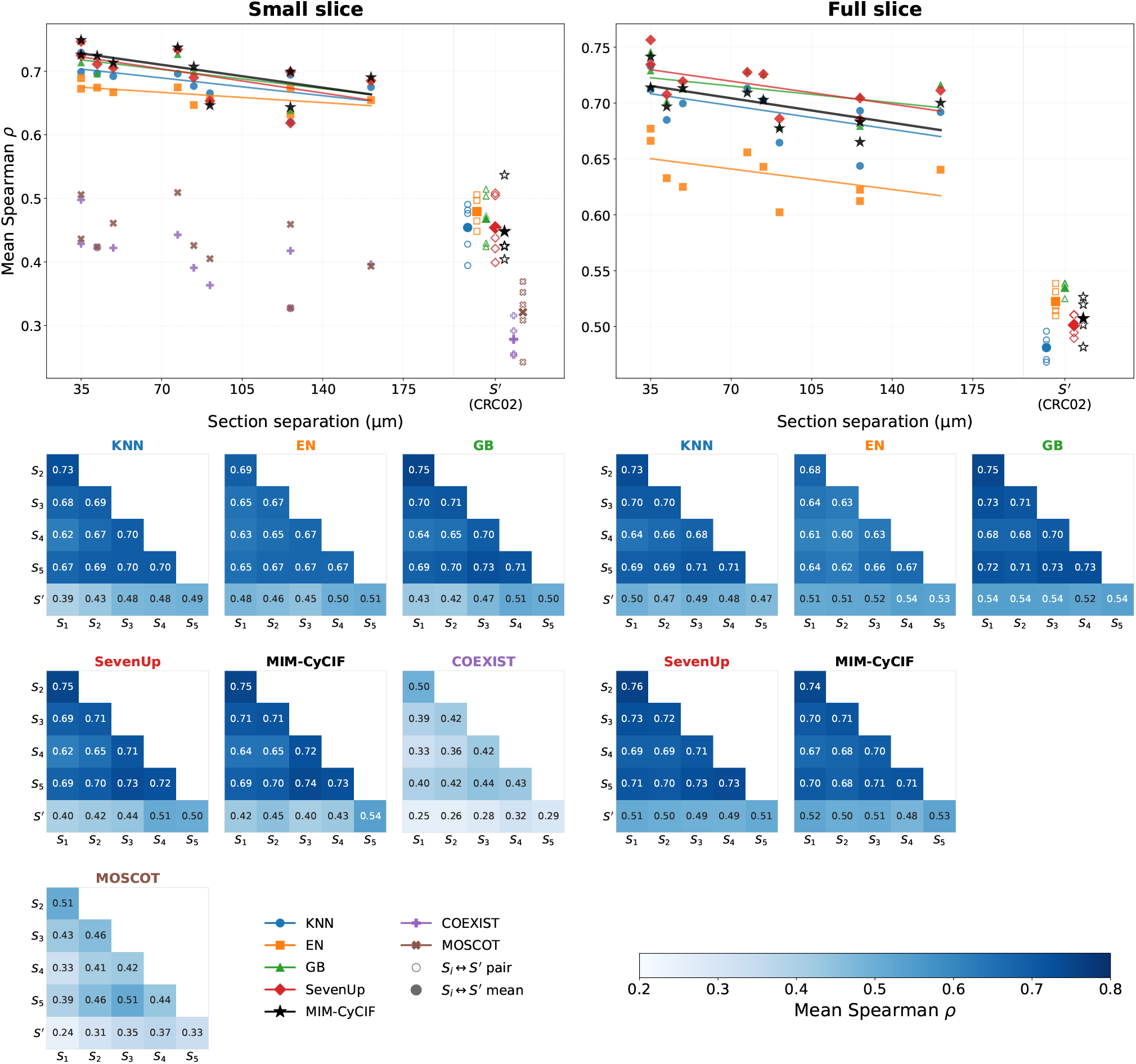
Mean Spearman *ρ* of proteins predicted in Benchmark 2 across different slice pairs.

The results show a *slight* negative correlation between inter-slice distance and prediction performance for all methods, and a *large* drop in performance when *S*^′^ is involved. Learning-based methods continue to dominate alignment-based methods. This allows us to draw the following insight: learning-based methods succeed in learning the *patient-specific* biological microenvironment (i.e., the relationship between known and unique proteins within a patient) from a single slice. This environment, clearly, does not change sufficiently across slices within a ≈ 100*µm* vicinity of each other to significantly affect cross-slice prediction accuracy. It does, however, change so between patients. Thus all of our learning-based methods have the following limitation: a method trained on one patient cannot reliably predict on another. Notably the alignment-based methods, besides still being the worst-performing overall, do not suffer as great of a cross-patient hit in performance. This is because they do not learn from (and thereby specialize to) a given patient before imputing on another. Their alignment-based imputation is thus shown to be more resilient to cross-patient imputation, although the performance remains relatively poor.

## 4 Discussion

We introduced and motivated the cross-slice protein imputation (CSPI) problem, and evaluated several methods for the solution of this problem. These methods can be broadly classified as learning-based or alignment-based, where a method of the former type will *learn* a relationship between common and unique proteins on one slice, to be used as a predictor on the other slice. A method of the latter type will instead *align* the cells and features of both slices to enable a transfer of unique protein expressions. For benchmarking, we addressed two major questions (i) How do the known protein sets in each slice affect the imputation performance across methods? (ii) How does choosing slices further apart in space affect imputation performance across methods? We selected these two questions to streamline design of spatial protein benchmarks where optimize selection of protein targets and tissue slices along the Z-axis of tissue blocks. These two parameters are critical because the optimization will allow to perform spatial proteomic profiling with minimal protein antibody sets and maximum slice distances ;. The resulting benchmark will inform on a diverse phenotypic coverage as well as a high tissue depth leading to comprehensive studies of tissue architecture, cell communication networks and therapeutically actionable multicellular mechanisms.

We have three major findings. First, for all benchmarks, learning-based methods significantly outperform alignment-based methods in protein imputation accuracy, measured both via Spearman and Pearson correlation. Second, across all methods, increasing both the size of the common protein set *P*_*c*_ and the number of cells per slice increases imputation accuracy. Third, cross-patient imputation is far less accurate than intra-patient imputation. In the latter case, there is a notable decrease in imputation accuracy as the slices are chosen further apart in space.

To monitor computational runtime for each benchmark and method, we recorded program execution time for all of our CSPI solution methods, for all of benchmark 1. With slices 1 and 2 fixed as *S*_1_ and *S*_2_, we observed how long each method took to impute the missing proteins across slices – this includes the time needed for learning/alignment – and how this time varies with different partitions of the protein space *P*, as well as with number of cells per slice. Table 5 shows the runtimes of all methods for the scaled down (10 000 cells) benchmark 1. For the the full-scale benchmark 1, these runtimes are shown in Table 6.

**Table 5:**
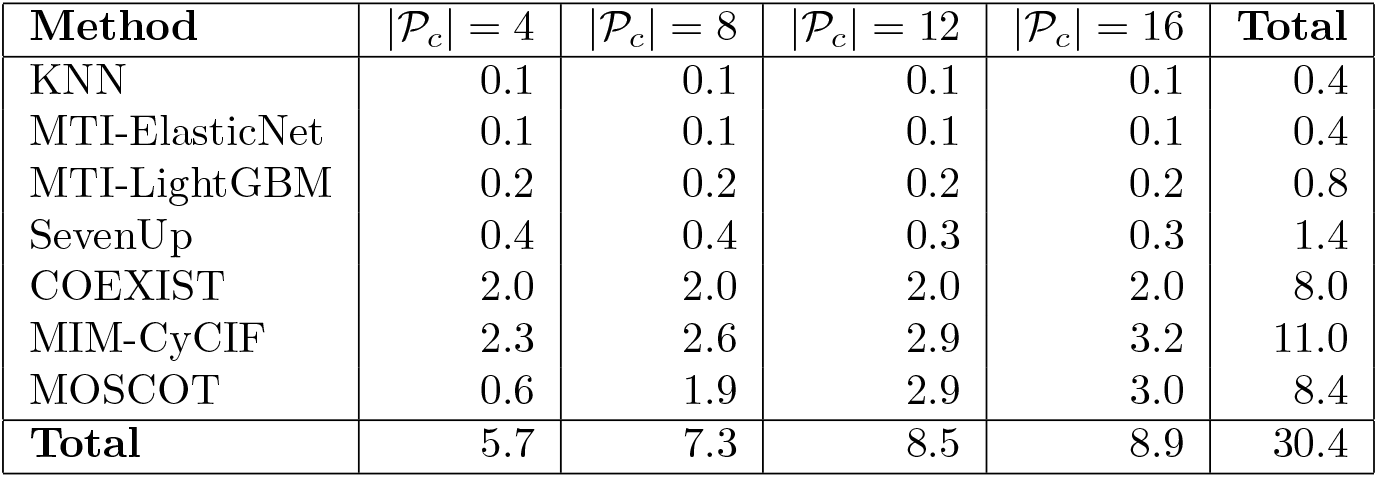
10,000-cell runtimes in minutes (*S*_1_ ↔ *S*_2_, fixed spatial center). COEXIST runtime is flat across panel sizes as it is dominated by the *N × N* cell-matching cost. MOSCOT and MIM-CyCIF both scale with |*P*_*c*_| : MOSCOT because the expression cost matrix grows with the shared panel, MIM-CyCIF because a larger input feature set requires more computation per forward pass.

| Method | $\vert \mathcal{P}_c\vert = 4$ | $\vert \mathcal{P}_c\vert = 8$ | $\vert \mathcal{P}_c\vert = 12$ | $\vert \mathcal{P}_c\vert = 16$ | Total |
| --- | --- | --- | --- | --- | --- |
| KNN | 0.1 | 0.1 | 0.1 | 0.1 | 0.4 |
| MTI-ElasticNet | 0.1 | 0.1 | 0.1 | 0.1 | 0.4 |
| MTI-LightGBM | 0.2 | 0.2 | 0.2 | 0.2 | 0.8 |
| SevenUp | 0.4 | 0.4 | 0.3 | 0.3 | 1.4 |
| COEXIST | 2.0 | 2.0 | 2.0 | 2.0 | 8.0 |
| MIM-CyCIF | 2.3 | 2.6 | 2.9 | 3.2 | 11.0 |
| MOSCOT | 0.6 | 1.9 | 2.9 | 3.0 | 8.4 |
| **Total** | 5.7 | 7.3 | 8.5 | 8.9 | 30.4 |

**Table 6:**
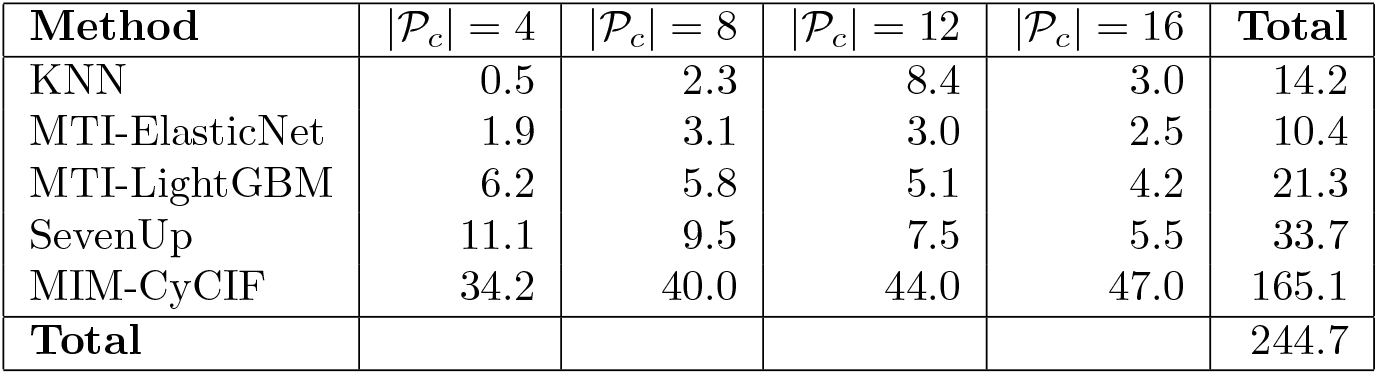
Full-slice (∼400k cells) runtimes in minutes. GPU–accelerated methods (MTI-LightGBM, SevenUp) run faster as |*P*_*c*_| grows because fewer target proteins require fitting. MIM-CyCIF trends the opposite way — a larger shared panel means more input features per forward pass.

| Method | $\vert \mathcal{P}_c\vert = 4$ | $\vert \mathcal{P}_c\vert = 8$ | $\vert \mathcal{P}_c\vert = 12$ | $\vert \mathcal{P}_c\vert = 16$ | Total |
| --- | --- | --- | --- | --- | --- |
| KNN | 0.5 | 2.3 | 8.4 | 3.0 | 14.2 |
| MTI-ElasticNet | 1.9 | 3.1 | 3.0 | 2.5 | 10.4 |
| MTI-LightGBM | 6.2 | 5.8 | 5.1 | 4.2 | 21.3 |
| SevenUp | 11.1 | 9.5 | 7.5 | 5.5 | 33.7 |
| MIM-CyCIF | 34.2 | 40.0 | 44.0 | 47.0 | 165.1 |
| **Total** |  |  |  |  | 244.7 |

Tables 5 and 6 show that, with the exception of MIM-CyCIF, methods’ runtime generally does not increase with |*P*_*c*_|; indeed, with SevenUp it actually decreases.

This first benchmark, testing for various protein set configurations, evaluated the performance of all methods for varying common protein sets *P*_*c*_ and, separately, for either 10 000 or ≈ 400 000 cells per slice. Thereby, we were able to see how methods perform both with varying known data (either in number of common proteins or number of cells per slice) and relatively to each other. A key observation is that alignment-based methods significantly underperform when compared to learning-based methods. Expression of any protein has a complex relationship between cell type, its other (known) protein expressions, and many unaccountable factors. Learning based methods attempt to recover this highly nonlinear relationship from knowledge of the known protein set *P*_*c*_ for a training set of cells – in this benchmark, either 10 000 or ≈ 400 000 cells. In contrast, alignment-based methods *transfer* a convex combination of protein expressions known in an adjacent slice based on a method-dependent computed ‘similarity score’ between cells in adjacent slices. Such linear-interpolative approaches fail to capture the complexity of the *P* → *P*_1 | 2_ relationships, as is reflected in the performance numbers.

Specifically, the results of Benchmark 1 show that MIM-CyCIF and SevenUp are the best performing methods regardless of how many proteins are shared between slices, with MIM performing better with less training data and SevenUp with more. MIM trains a single FT-transformer to perform simultaneous prediction of all unknown proteins – this is beneficial for learning complex joint relationships for small training data, but the complexity of this approach makes it vulnerable to noisy data and computationally intensive to train for larger datasets. SevenUp, however, trains one LightGBM model per target protein – this approach benefits from and scales well with more training data and more common proteins. Thus, practitioners could lean towards MIM in cases where the slices have few cells and few common proteins (such as in tissue microarrays), and towards SevenUp’s architecture where more data is available.

For the second benchmark, where the distances between slice were varied, the main result is that the distance between intra-patient slices has a far smaller effect on imputation accuracy than initially expected, and that performance really suffers when imputing cross-patient. For alignment-based methods this is naturally the case – the spatial alignment of the two slices which they take as a prerequisite cannot be well-guaranteed in this setting. For learning-based methods, this result shows that the *P* _*c*_ → *P*_1| 2_ relationship learned on one slice in one patient is of limited use for imputation in another patient. This reflects cross-patient differences in tissue microenvironment, which a model trained on a single patient cannot be expected to account for, and suggests further exploration in models trained across multiple patients.

The conclusions of benchmark 2 are consistent with those of benchmark 1, where both MIM and 7-UP are the best performers over all slice pairings. Perhaps unsurprisingly, we see performance of all methods drop as slices are chosen to be further apart. This performance drop is very significant when they are chosen from different patients. Thus the methods we evaluate here are best trained and deployed on a per-patient scale, with generalization across patients remaining a subject of future work.

Finally, the alignment-based methods, originally developed for simpler cases of feature and cell transfer (MOSCOT) and for identifying cells common to spatially-adjacent slices (COEXIST), are found to be less performant in the CSPI problem when compared to learning-based methods. Despite this, however, their performance is not wholly inadequate on its own, which leads us to recommend the use of such methods for alignment of *spatial* features as opposed to protein expressions. Such alignments can be used to recover inter-slice tissue structure where none is known. See Appendix A for more details.

## Code Availability

The code and instructions to reproduce the results of this paper can be found at https://github.com/keshakot/cross_slice_protein_imputation.

## A Application to 3D Tissue Reconstruction

The previous benchmarks have shown a consistent pattern – alignment-based methods perform significantly worse at solving the CSPI problem than learning-based methods. In this section we provide an alternative application of alignment-based methods to the task of 3D tissue reconstruction, which naturally complements the CSPI problem and its motivation.

The 3D reconstruction problem is the problem of inter-slice interpolation – if *S*_1_ is at *z* = 0 and *S*_2_ is at *z* = 1, what is at *z* ∈ (0, 1)? The *what* refers to both the positions of individual cells as well as the protein expressions thereof.

Alignment-based methods are a natural fit for 3D reconstruction, since they provide a mapping/association between cells in adjacent slices. Once this mapping is in place, it can be used to transfer any features between cells in adjacent slices. One way to visualize this is to consider the mapping as a matrix 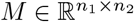, where *n*_*i*_ is the number of cells in slice *i*. The entry [*M*]_*i,j*_ is the strength of association of cell *i* in slice 1 to cell *j* in slice 2. Given a feature vector 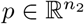 representing the expression intensities of a protein in slice 2, we can make transfer this vector to slice 1 as follows:

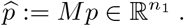

Here, the vector 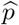 is the prediction of the feature’s expression in slice 1. Conversely, features in slice 1 can be transferred to slice 2 by left-multiplication by *M* ^⊤^. As our previous benchmarks have shown, prediction of protein expressions by such methods has low accuracy when compared to learning-based methods. However, such methods allow for a natural form of interpolation between the two slices. For instance, denote the (*x, y*) positions of cells in slice 1 as 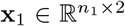, and similarly **x**_2_ for slice 2. We can learn an alignment *M* such that

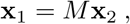

i.e. this alignment respects the *known* positions of cells within the slices. Now, to predict the positions of cells between the slices, let *θ* (0, 1). Then with slice 1 at *z* = 0 and slice 2 at *z* = 1, we can predict the distribution of cells at the slice at *z* = *θ* as the *n*_1_*n*_2_ locations (1 − *θ*)[***x***_1_]_*i*_ + *θ*[***x***_2_]_*j*_ for all (*i, j*) ∈ ⟬*n*_1_ *× n*_2_ ⟭, each with the associated intensity [*M*]_*i,j*_. This can then be downsampled via an appropriate scheme (e.g. KNN, KDE fitting) to obtain a distribution with the desired number of cells, usually ≈ *n*_1|2_.

To demonstrate this approach, we solve a fused Gromov-Wasserstein (FGW) problem (Titouan et al., 2019) via MOSCOT between slices *S*_1_ ↔ *S*_3_ and *S*_3_ ↔ *S*_5_ as in Table 2, and use the solutions to predict the cell positions in slices *S*_2_ and *S*_4_. We also perform a refinement (shown as OT-TPS in the figure) where cells are forced to be transported bijectively between the slices, meaning no cell mass is split. This results in a more ‘crisp’, structure-preserving reconstruction than pure FGW. To provide a baseline, we also perform naive pooling, which simply takes 50% of cells from each slice to form the predicted intermediate slice. We show the results in Figure 5.

**Figure 5:**
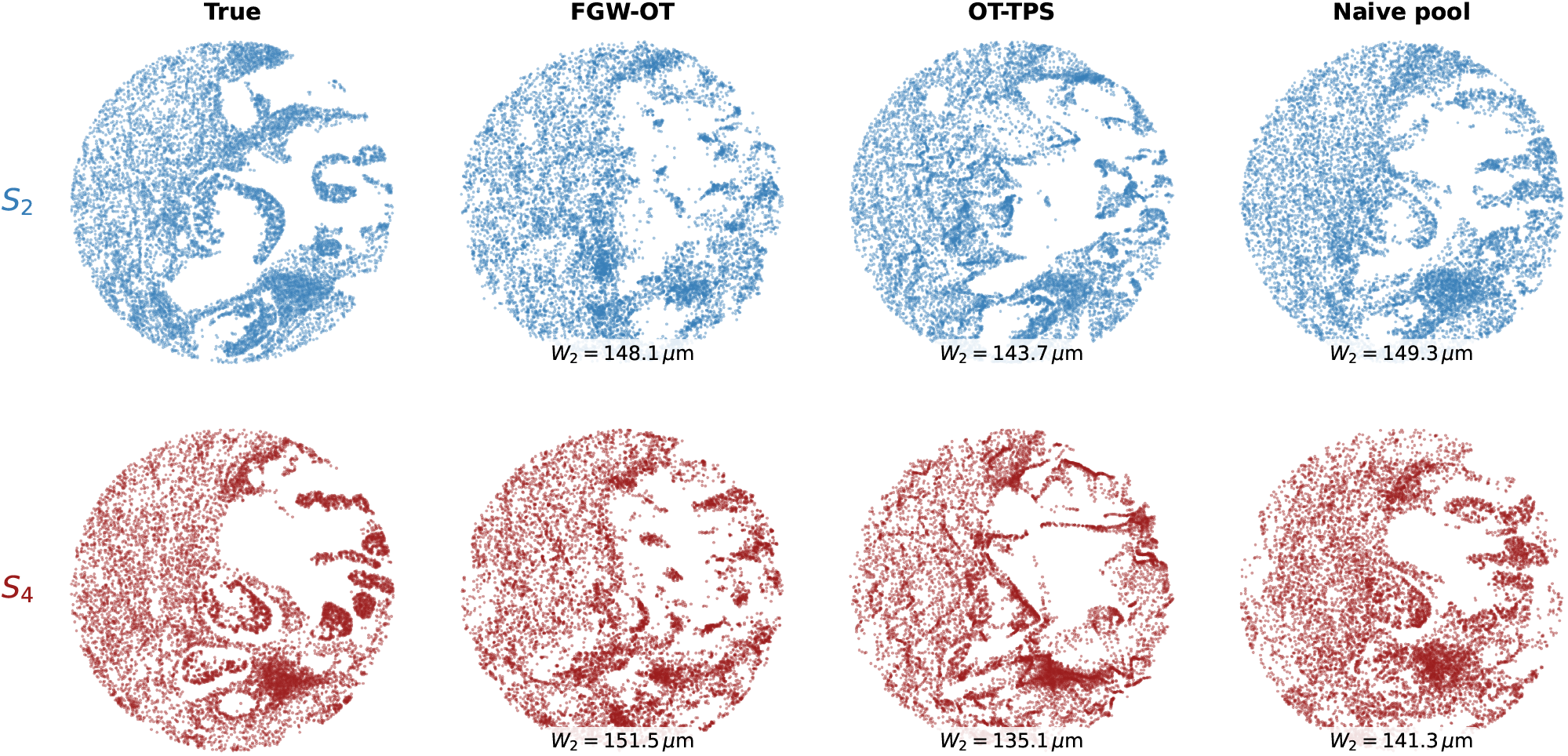
Known vs. reconstructed slices *S*_2_ and *S*_4_ as in Table 2, done with FGW, both direct and TPS), and naive pooling. Wasserstein *W*_2_ distances are reported between each prediction and the ground truth (True) slice.

As evident from Figure 5, the general predicted structure of the slices matches closely with that of the known slices, for all methods. The FGW interpolation tends to smoothen the details of both endpoint slices, thus resulting in loss of small-scale feature fidelity. The OT-TPS bijection counteracts this, and indeed this variant has the lowest Wasserstein *W*_2_ distance (Gabriel & Marco, 2019, Sec. 2.4) between the prediction and the ground-truth slice, both for *S*_2_ and *S*_4_. The use of alignment-based methods for 3D reconstruction is a potentially powerful technique for prediction of tissue structure where none is available.

## Footnotes

1 In practice, these expressions are given in tabular format. For instance, 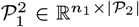 is a table of *n*_1_ rows, where row *i* gives the expression of proteins _2_ for cell *i* of the *n*_1_ cells in slice 1. Similarly, we understand 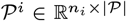 to be the full table of protein expressions for slice *i*.

2 Both the number of cells (10 000) and the reference point for this scaled-down benchmark were chosen arbitrarily to reduce computation time while providing enough training data for learning-based methods, along with nontrivial tissue structure.

3 Scaling to this number of cells *is* feasible, but requires considerable runtime and lower-rank approximations of alignments. For instance, a naive alignment between full slices requires a matrix of ≈ 400 000^2^ entries.

## References

Alina Bollhagen and Bernd Bodenmiller. Highly multiplexed tissue imaging in precision oncology and translational cancer research. 14(11):2071–2088. ISSN 2159-8274. doi: 10.1158/2159-8290.CD-23-1165. URL https://www.ncbi.nlm.nih.gov/pmc/articles/PMC11528208/.

Jacob Devlin, Ming-Wei Chang, Kenton Lee, and Kristina Toutanova. BERT: Pre-training of deep bidirectional transformers for language understanding. In Proceedings of the 2019 Conference of the North American Chapter of the Association for Computational Linguistics: Human Language Technologies (NAACLHLT), pp. 4171–4186, 2019. doi: 10.18653/v1/N19-1423.

Ofer Elhanani, Raz Ben-Uri, and Leeat Keren. Spatial profiling technologies illuminate the tumor microenvironment. 41(3):404–420. ISSN 1535-6108, 1878-3686. doi: 10.1016/j.ccell.2023.01.010. URL https://www.cell.com/cancer-cell/abstract/S1535-6108(23)00010-7.

Peyré Gabriel and Cuturi Marco. Computational optimal transport with applications to data sciences. Foundations and trends® in machine learning, 11(5-6):355–607, 2019.

Jérôme Galon, Anne Costes, Fatima Sanchez-Cabo, Amos Kirilovsky, Bernhard Mlecnik, Christine Lagorce-Pagès, Marie Tosolini, Matthieu Camus, Anne Berger, Philippe Wind, Franck Zinzindohoué, Patrick Bruneval, Paul-Henri Cugnenc, Zlatko Trajanoski, Wolf-Herman Fridman, and Franck Pagès. Type, density, and location of immune cells within human colorectal tumors predict clinical outcome. 313(5795):1960–1964. ISSN 1095-9203. doi: 10.1126/science.1129139.

Charlotte Giesen, Hao A. O. Wang, Denis Schapiro, Nevena Zivanovic, Andrea Jacobs, Bodo Hattendorf, Peter J. Schüffler, Daniel Grolimund, Joachim M. Buhmann, Simone Brandt, Zsuzsanna Varga, Peter J. Wild, Detlef Günther, and Bernd Bodenmiller. Highly multiplexed imaging of tumor tissues with subcellular resolution by mass cytometry. 11(4). ISSN 1548-7105. doi: 10.1038/nmeth.2869.

Yury Goltsev, Nikolay Samusik, Julia Kennedy-Darling, Salil Bhate, Matthew Hale, Gustavo Vazquez, Sarah Black, and Garry P. Nolan. Deep profiling of mouse splenic architecture with CODEX multiplexed imaging. 174(4):968–981.e15. ISSN 0092-8674, 1097-4172. doi: 10.1016/j.cell.2018.07.010. URL https://www.cell.com/cell/abstract/S0092-8674(18)30904-8.

Yury Gorishniy, Ivan Rubachev, Valentin Khrulkov, and Artem Babenko. Revisiting deep learning models for tabular data. In Advances in Neural Information Processing Systems, volume 34, pp. 18932–18943. Curran Associates, Inc., 2021.

Charles R. Harris, K. Jarrod Millman, Stéfan J. van der Walt, Ralf Gommers, Pauli Virtanen, David Cournapeau, Eric Wieser, Julian Taylor, Sebastian Berg, Nathaniel J. Smith, Robert Kern, Matti Picus, Stephan Hoyer, Marten H. van Kerkwijk, Matthew Brett, Allan Haldane, Jaime Fernández del Río, Mark Wiebe, Pearu Peterson, Pierre Gérard-Marchant, Kevin Sheppard, Tyler Reddy, Warren Weckesser, Hameer Abbasi, Christoph Gohlke, and Travis E. Oliphant. Array programming with NumPy. Nature, 585(7825):357–362, 2020. doi: 10.1038/s41586-020-2649-2.

Robert T Heussner, Cameron F Watson, Christopher Z Eddy, Kunlun Wang, Eric M Cramer, Allison L Creason, Gordon B Mills, and Young Hwan Chang. Coexist: Coordinated single-cell integration of serial multiplexed tissue images. PLoS computational biology, 21(8):e1013325, 2025.

John W. Hickey, Yuqi Tan, Garry P. Nolan, and Yury Goltsev. Strategies for accurate cell type identification in CODEX multiplexed imaging data. Frontiers in Immunology, 12:727626, 2021. doi: 10.3389/fimmu.2021.727626.

Leeat Keren, Marc Bosse, Diana Marquez, Roshan Angoshtari, Samir Jain, Sushama Varma, Soo-Ryum Yang, Allison Kurian, David Van Valen, Robert West, Sean C. Bendall, and Michael Angelo. A structured tumor-immune microenvironment in triple negative breast cancer revealed by multiplexed ion beam imaging. 174(6):1373–1387.e19. ISSN 1097-4172. doi: 10.1016/j.cell.2018.08.039.

Raphael Kirchgaessner, Cameron Watson, Allison Creason, Kaya Keutler, and Jeremy Goecks. Imputing single-cell protein abundance in multiplex tissue imaging. Nature Communications, 16(1):4747, 2025.

Dominik Klein, Giovanni Palla, Marius Lange, Michal Klein, Zoe Piran, Manuel Gander, Laetitia Meng-Papaxanthos, Michael Sterr, Aimee Bastidas-Ponce, Marta Tarquis-Medina, et al. Mapping cells through time and space with moscot. biorxiv. 2023, 2005(2011):540374, 2023.

Jia-Ren Lin, Benjamin Izar, Shu Wang, Clarence Yapp, Shaolin Mei, Parin M. Shah, Sandro Santagata, and Peter K. Sorger. Highly multiplexed immunofluorescence imaging of human tissues and tumors using t-CyCIF and conventional optical microscopes. Elife, 7:e31657, July 2018. ISSN 2050-084X. doi: 10.7554/eLife.31657.

Jia-Ren Lin, Shu Wang, Shannon Coy, Yu-An Chen, Clarence Yapp, Madison Tyler, Maulik K Nariya, Cody N Heiser, Ken S Lau, Sandro Santagata, et al. Multiplexed 3d atlas of state transitions and immune interaction in colorectal cancer. Cell, 186(2):363–381, 2023.

Ilya Loshchilov and Frank Hutter. SGDR: Stochastic gradient descent with warm restarts. In International Conference on Learning Representations (ICLR), 2017.

Ilya Loshchilov and Frank Hutter. Decoupled weight decay regularization. In International Conference on Learning Representations (ICLR), 2019.

Jeffrey R. Moffitt, Emma Lundberg, and Holger Heyn. The emerging landscape of spatial profiling technologies. 23(12):741–759. ISSN 1471-0064. doi: 10.1038/s41576-022-00515-3. URL https://www.nature.com/articles/s41576-022-00515-3.

Fabian Pedregosa, Gaël Varoquaux, Alexandre Gramfort, Vincent Michel, Bertrand Thirion, Olivier Grisel, Mathieu Blondel, Andreas Müller, Joel Nothman, Gilles Louppe, Peter Prettenhofer, Ron Weiss, Vincent Dubourg, Jake Vanderplas, Alexandre Passos, David Cournapeau, Matthieu Brucher, Matthieu Perrot, and Édouard Duchesnay. Scikit-learn: Machine Learning in Python. Journal of Machine Learning Research, 12:2825–2830, October 2011. doi: 10.48550/arXiv.1201.0490.

Christian M. Schürch, Salil S. Bhate, Graham L. Barlow, Darci J. Phillips, Luca Noti, Inti Zlobec, Pauline Chu, Sarah Black, Janos Demeter, David R. McIlwain, Shigemi Kinoshita, Nikolay Samusik, Yury Goltsev, and Garry P. Nolan. Coordinated cellular neighborhoods orchestrate antitumoral immunity at the colorectal cancer invasive front. 182(5):1341–1359.e19. ISSN 0092-8674. doi: 10.1016/j.cell.2020.07.005. URL https://www.sciencedirect.com/science/article/pii/S0092867420308709.

Zachary Sims, Gordon B Mills, and Young Hwan Chang. Mim-cycif: masked imaging modeling for enhancing cyclic immunofluorescence (cycif) with panel reduction and imputation. Communications Biology, 7(1):409, 2024.

Vayer Titouan, Nicolas Courty, Romain Tavenard, Chapel Laetitia, and Rémi Flamary. Optimal transport for structured data with application on graphs. In Kamalika Chaudhuri and Ruslan Salakhutdinov (eds.), Proceedings of the 36th International Conference on Machine Learning, volume 97 of Proceedings of Machine Learning Research, pp. 6275–6284. PMLR, 09–15 Jun 2019. URL https://proceedings.mlr.press/v97/titouan19a.html.

Eric Wu, Alexandro E Trevino, Zhenqin Wu, Kyle Swanson, Honesty J Kim, H Blaize D’Angio, Ryan Preska, Aaron E Chiou, Gregory W Charville, Piero Dalerba, et al. 7-up: Generating in silico codex from a small set of immunofluorescence markers. PNAS nexus, 2(6):pgad171, 2023.

